# Effects of Starter Feeds of Different Physical Forms on Rumen Fermentation and Microbial Composition for Pre-weaning and Post-weaning Lambs

**DOI:** 10.1101/2020.08.03.235580

**Authors:** Yong Li, Yanli Guo, Chengxin Zhang, Xiaofang Cai, Peng Liu, Cailian Li

## Abstract

This study aimed to evaluate the effects of starter feeds of different physical forms on rumen fermentation and microbial composition for lambs. Twenty-four eight-day-old male Hu lambs (5.04 ± 0.75 kg body weight) were fed either milk replacer (MR) and pelleted starter feed (PS), or MR and textured starter feed (TS) in pre-weaning (day 8 to 35) and post-weaning (day 36 to 42) lambs. And the MR was fed by bottles to lambs at 2% of body weight at day 8 divided as three equal amounts at 08:00, 14:00 and 20:00 in pre-weaning. And the lambs were readily availed starter feeds and clean fresh water in the whole experiment. Six lambs for each treatment were euthanized at day 21 or 42 for sampling. The results showed the total volatile fatty acids, propionate and butyrate of rumen liquid in TS groups were all higher than them in PS groups respectively for pre-weaning and post-weaning lambs (*P* < 0.05), and the pH of rumen liquid in TS group was lower than it in PS group for post-weaning lambs (*P* < 0.05). Moreover, the pH of rumen and OTUs in TS group had trends to lower than them in PS group for pre-weaned lambs (*P* = 0.061, *P* = 0.066). TS established the predominant Phylum, *Bacteroidetes*, earlier than PS, and increased significantly the relative abundances of *Sharpea* compared to PS at level of genus (*P* < 0.05) for pre-weaning and post-weaning lambs. TS were more benefits to trigger rumen development for lambs.

**IMPORTANCE:** Early use of starter feed could trigger rumen fermentation and establishment of dominant flora, which were in favour of growth and development of rumen for ruminants. The physical form of starter feed is one of the important factors to promote rumen fermentation and establishment of dominant flora for ruminants of transition. However, limited study on effects of physical forms of starter feeds, especially the texturized starters containing steam-flaked grains, to rumen fermentative pattern and microbial composition for pre-weaning and post-weaning lambs to date. It was necessary to investigate the effects of physical form of starter feed on rumen fermentation and microbial composition for lambs. The significance of our research showed TS were better benefits to promote the rumen fermentation and establishment of dominant flora for lambs, which will greatly enhance our understanding of physical forms of starter feeds, leading to broader studies on rumen development for lambs.

## INTRODUCTION

It is commonly known that a lot of complicated and diverse microbe, such as protozoan, bacteria, archeobacteria and fungi, existed in rumen of ruminant, whose interaction played an important role to maintain stable environments of rumen and health of animals (1). However, at birth, young ruminants did not possess anaerobic microbial population in their rumens. And establishment of rumen microbiota was very necessary to the physiological development of the rumens and the animal’s abilities to convert plant feed into products that can be utilized by the animal for maintenance and production (2). Previous studies showed the introduction of solid diet around weaning could promote the establishment of anaerobic microbial ecosystem and formation of fermentation processes, which were benefits to trigger the growth and development of the rumen (3).

Furthermore, processing of grain and ingredients of starter feeds and chemical and structural composition in dietary could affect fermentative pattern of rumen (4, 5). Feeding starters containing fine particles in form of mash or processed in form of pelleted could trigger rapid ruminal acid production from fermentation of carbohydrates (6), decreased ruminal pH (7). Consumption of starter feed containing highly fermentable carbohydrate by lambs or calves could increase the ruminal concentrations of volatile fatty acids (VFAs), particularly propionate and butyrate (8, 9), which was essential chemical stimulations for the rapid development of rumen epithelium (10). Furthermore, consumption of solid feed could promote the acquisition of anaerobic microbes and establishment of rumen fermentation (11).

Additionally, microbial colonization can also affect rumen development and function during early life, and the diversities and function of rumen microbe during early life was given concerned widely in recent years and next-generation sequencing was widely used to study rumen microbial ecology. Jami et al. and Li et al. found that the rumen microbe of prior to weaning calves has a similar functional capacity as that of a mature ruminant using next generation DNA (2, 12). Chen et al. studied the changes in bacterial diversity associated with epithelial tissue in the beef cow rumen during the transition to a high-grain diet (13). Jiao et al. taxonomic identified the ruminal epithelial bacterial diversity during rumen development in Goats (14). However, limited information is available on how changes in the physical forms of starter feeds, especially the texturized starters containing steam-flaked grains, influence the rumen fermentative pattern and microbial composition for pre-weaning and post-weaning lambs to date. And there was not sufficient research on diversities and function of rumen microbe of starter feeds of different physical forms used next generation DNA for pre-weaning and post-weaning lambs.

Hence, the objectives of experiment were to elaborate the relation of effect of starter feeds of different physical forms on rumen fermentation and microbial composition for pre-weaning and post-weaning lambs by diversities and functions of rumen microbe (next-generation sequencing, 16S rDNA gene sequencing) so as to explain the reasons that effects of starter feeds of different physical forms on growth and development of rumen for pre-weaning and post-weaning lambs to certain extent.

## RESULTS AND ANALYSIS

### Rumen fermentative parameters

The total VFAs, propionate and butyrate of rumen in TS groups were all higher than them in PS groups respectively for pre-weaned and post-weaned lambs (*P* < 0.05), and the pH of rumen in TS group was lower than it in PS group for post-weaned lambs (*P* < 0.05, Table 1). Moreover, the pH of rumen in TS group had trends to lower than it in PS group in pre-weaned lambs (*P* = 0.061).

**Table 1.** Rumen fermentative parameters of lambs.

| Items | Pelleted | Textured | SEM | *P-value* |
| --- | --- | --- | --- | --- |
| Pre-weaning |  |  |  |  |
| pH | 5.96 | 5.60 | 0.12 | 0.061 |
| Total volatile fatty acids (mmol/L) | 24.41 | 30.04 | 1.72 | 0.046 |
| ) |  |  |  |  |
| Acetate (mmol/L) | 13.94 | 15.56 | 1.30 | 0.414 |
| Propionate (mmol/L) | 6.69 | 9.07 | 0.72 | 0.046 |
| Butyrate (mmol/L) | 2.02 | 3.16 | 0.19 | 0.006 |
| Isobutyrate (mmol/L) | 0.34 | 0.48 | 0.05 | 0.110 |
| Valerate (mmol/L) | 0.85 | 1.08 | 0.21 | 0.465 |
| Isovalerate (mmol/L) | 0.56 | 0.70 | 0.09 | 0.320 |
| Acetate to propionate | 2.16 | 1.76 | 0.26 | 0.308 |
| Ammonia nitrogen (mg/100ml) | 12.51 | 14.87 | 0.93 | 0.107 |
| Post-weaning |  |  |  |  |
| pH | 5.48 | 5.14 | 0.07 | 0.011 |
| Total volatile fatty acids (mmol/L) | 65.59 | 70.72 | 1.51 | 0.046 |
| ) |  |  |  |  |
| Acetate (mmol/L) | 38.89 | 38.02 | 1.65 | 0.739 |
| Propionate (mmol/L) | 16.73 | 21.56 | 1.49 | 0.044 |
| Butyrate (mmol/L) | 5.79 | 7.67 | 0.50 | 0.023 |
| Isobutyrate (mmol/L) | 0.45 | 0.46 | 0.04 | 0.989 |
| Valerate (mmol/L) | 3.03 | 2.20 | 0.58 | 0.332 |
| Isovalerate (mmol/L) | 0.70 | 0.81 | 0.08 | 0.317 |
| Acetate to propionate | 2.52 | 1.81 | 0.29 | 0.153 |
| Ammonia nitrogen (mg/100ml) | 12.25 | 12.75 | 0.82 | 0.677 |

### Sequence qualities of 16S rRNA genes and alpha diversities of rumen microbe

Raw sequences were joined together, optimized, quality controlled, then 1,648,195 high quality sequences were obtained. Every sample obtained 82,410↑ V3-V4 16S r RNA effective tags averagely. The length of effective tags was between 416 bp-426 bp. The 2,248 OTUs were obtained in all, and every sample had 562 OTUs averagely (Fig 1). Rarefaction curve of 16S rRNA gene showed Good’ coverage was higher than 0.99. Based on similarity principles of 97% sequences between reads, OTUs coverage of sequences was adequate. As showed Fig 2, **the** rarefaction curve of 16S rRNA gene had trends to smooth, which showed sequences were reasonable. The span of rank abundance of 16S rRNA gene in the horizontal direction had trends to increase, which showed the abundance of species had trend to increase, and the span of rank abundance of 16S rRNA gene in the vertical direction had trends to smooth, which showed the distribution of species had trend to even. In a word, the sequences could reflect accurately the rumen microbial composition for lambs.

**Fig 1.**
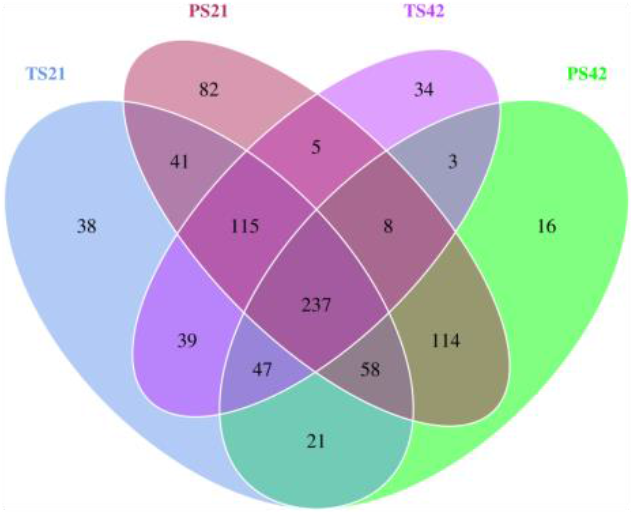
Venn graph

**Fig 2.**
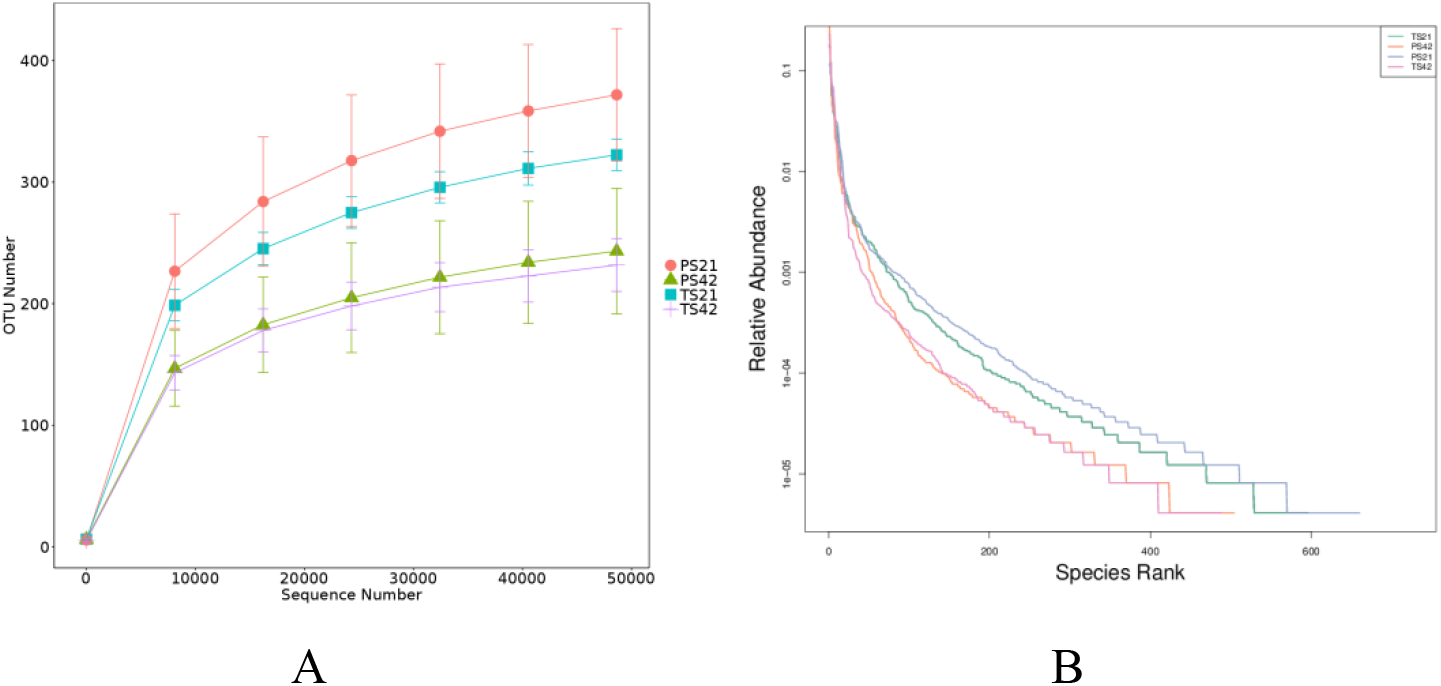
Rarefaction curve and Rank abundance of 16S rRNA gene Note: A, rarefaction curve; B, rank abundance.

Except for OTUs of PS had trend to higher than them of TS for pre-weaning lambs (*P* = 0.066), the physical form of starter feed did not affect the alpha diversities of rumen bacterial communities for pre-weaning and post-weaning lambs (*P* > 0.05, Table 2).

**Table 2.** Alpha diversities of rumen bacterial communities.

| Items | Pelleted | Textured | SEM | *P-value* |
| --- | --- | --- | --- | --- |
| Pre-weaning |  |  |  |  |
| OTU | 415.80 | 352.20 | 40.79 | 0.066 |
| ACE | 421.40 | 374.85 | 44.74 | 0.146 |
| Chao1 | 415.79 | 406.93 | 94.83 | 0.893 |
| Shannon | 3.74 | 4.09 | 0.27 | 0.407 |
| Simpson | 0.78 | 0.84 | 0.09 | 0.323 |
| Post-weaning |  |  |  |  |
| OTU | 279.00 | 258.40 | 48.90 | 0.557 |
| ACE | 282.18 | 263.49 | 43.66 | 0.556 |
| Chao1 | 272.76 | 258.52 | 40.10 | 0.645 |
| Shannon | 3.61 | 3.44 | 0.57 | 0.662 |
| Simpson | 0.81 | 0.79 | 0.09 | 0.766 |

### Beta diversities of rumen microbe

Principal co-ordinates analysis (PCoA) of rumen bacterial OTUs showed contribution rates of PC1 and PC2 to differences among samples were 21.80% and 19.69% respectively, which could reflect adequately the differences among samples. PCoA results showed differences of samples were minute in same groups (Fig 3). Non-metric multi-dimensional scaling analysis (NMDS) of rumen bacterial OTUs (Fig 4) showed stress value lower 0.2 (0.133), which could indicate accurately the data and reflect the significant differences of rumen microbial structure and diversity (*P* < 0.05).

**Fig 3.**
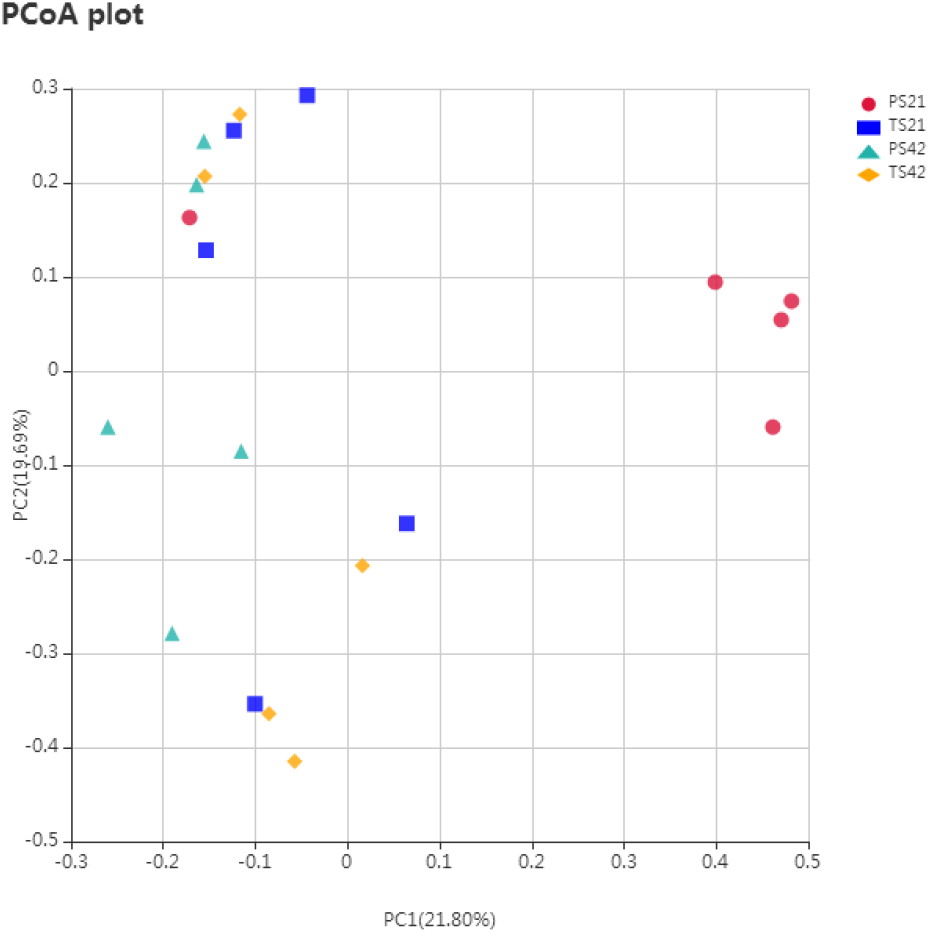
Principal co-ordinates analysis (PCoA) of rumen bacterial OTUs

**Fig 4.**
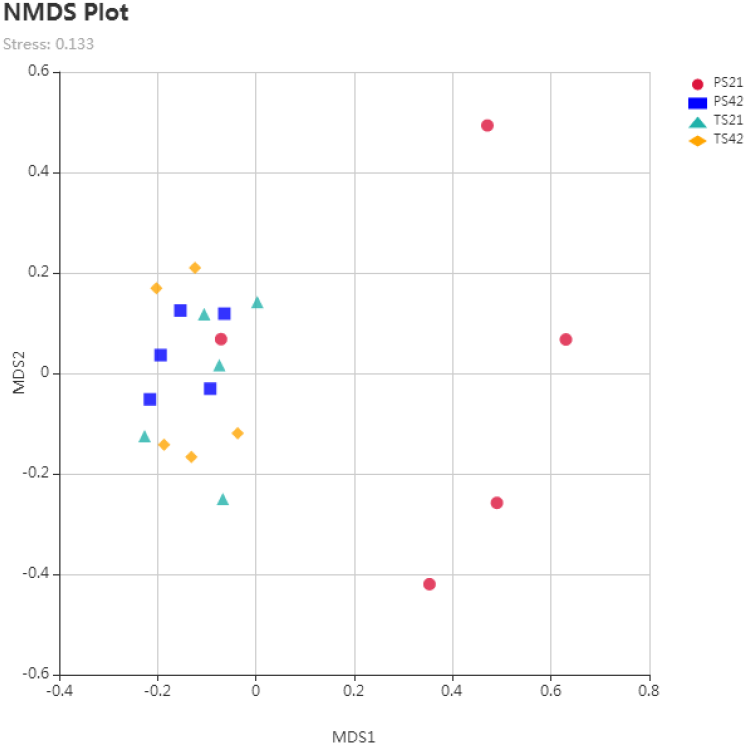
Non-metric multi-dimensional scaling analysis (NMDS) of rumen bacterial OTUs

### Effects of starter feeds of different physical forms on rumen microbe

Among the Phylum of top 10, the predominant Phylum was all *Bacteroidetes* and *Firmicutes* in every group, and their relative abundances were all higher than 24% (Fig 5). Only the relative abundances of *Bacteroidetes* of TS had trends to higher than them of PS for pre-weaning lambs (*P*=0.084). However, with intake of starter feed, the relative abundances of *Bacteroidetes* increased, and the relative abundances of *Firmicutes* decreased. The relative abundances of *Bacteroidetes* (61.96%) of TS had exceed them of *Firmicutes* (32.08%) and *Proteobacteria* (3.99%) for pre-weaning lambs, and become the first predominant Phylum, which were similar to them *Bacteroidetes* (65.36%) of TS for post-weaning lambs. But the relative abundances of *Bacteroidetes* (57.28%) of PS exceeded them of *Firmicutes* (31.21%) and *Proteobacteria* (1.15%) for post-weaning lambs, which were still lower than them of *Bacteroidetes* of TS for pre-weaning (61.96%) and post-weaning (65.36%) lambs.

**Fig 5.**
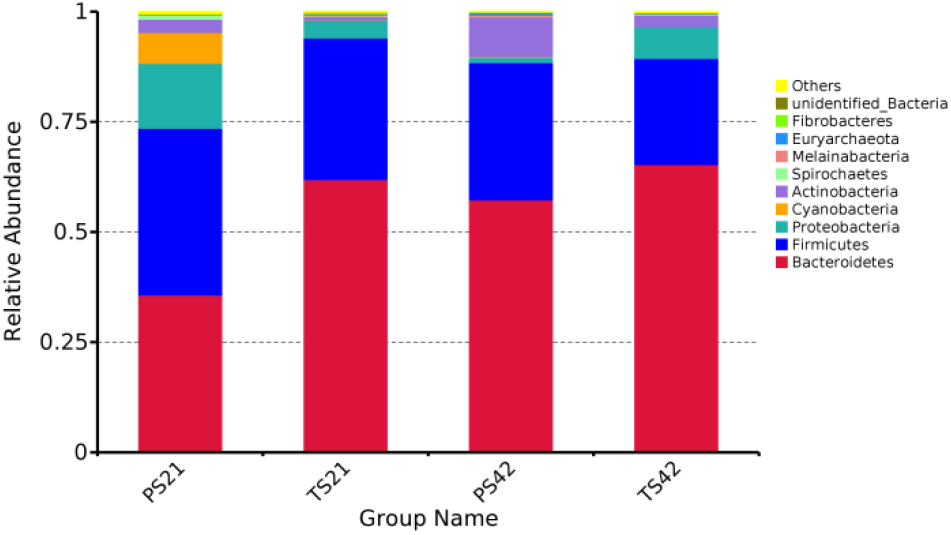
Phylum of top 10 of rumen microbe for lambs

It showed 153 rumen microflora were detected at the genus level between TS and PS for pre-weaning lambs, and the relative abundances of 13 rumen microflora were higher than 0.5%. The predominant microflora of PS and TS commonly were *unidentified_Prevotellaceae* (6.49% and 21.85%). The predominant microflora of PS peculiarly was *Lactobacillus* (14.53%), *Succinivibrio* (10.82%) and *unidentified_Cyanobacteria* (6.96%), and the predominant microflora of TS peculiarly was *Sharpea* (4.41%), *Dialister* (3.85%) and *Succinivibrio* (3.18%). The results of rumen microbe showed the PS increased significantly *unidentified_Clostridiales*, *Lactpcoccus*, *Sarcina*, *unidentified_Cyanobacteria*, and TS increased significantly *sharpea* and *Oribacterium* compared between PS and TS for pre-weaning lambs at the genus level (Fig 6, LDA>4).

**Fig 6.**
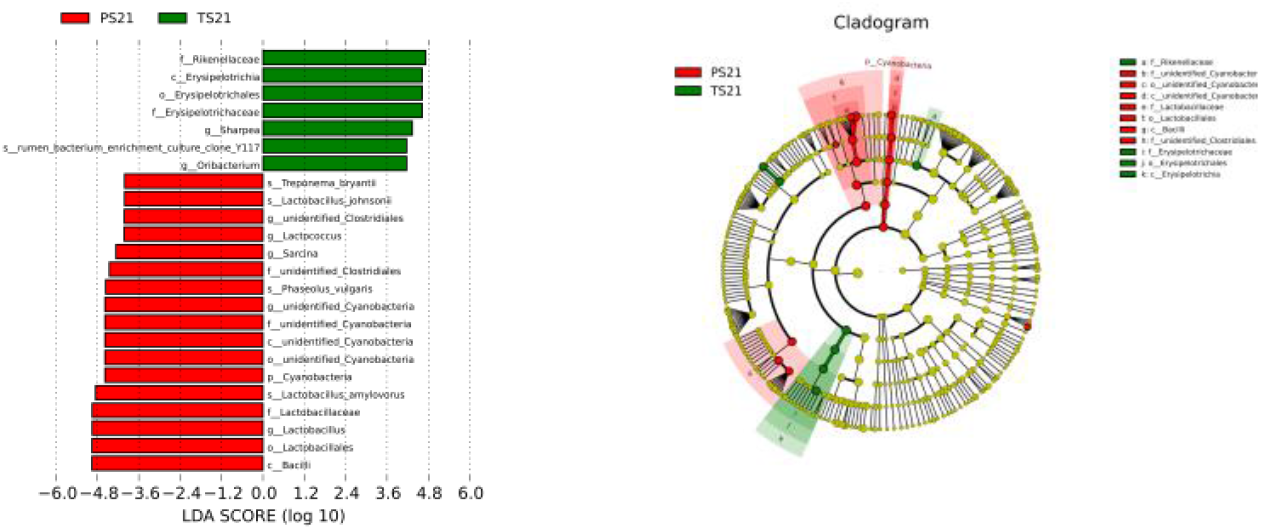
Value of linear discriminant analysis (LDA) effect size (LEf Se) on rumen microflora between two starter feeds for pre-weaned lambs

However, the effects of starter feed of two physical forms on rumen microflora were different for post-weaning lambs. And 139 rumen microflora were detected at the genus level between TS and PS for post-weaning lambs, and the relative abundances of 13 rumen microflora were higher than 0.5%. The predominant microflora of PS and TS commonly were *unidentified_Prevotellaceae* (29.57% and 38.49%). The predominant microflora of PS peculiarly was *Dialister* (7.23%), *unidentified_Lachnospiraceae* (6.72%), and the predominant microflora of TS peculiarly were *Sharpea* (7.43%) and *Succinivibrio* (6.87%). The results of rumen microbe showed only the TS increased significantly *sharpea* compared between PS and TS for post-weaned lambs at the genus level (Fig 7, LDA>4).

**Fig 7.**
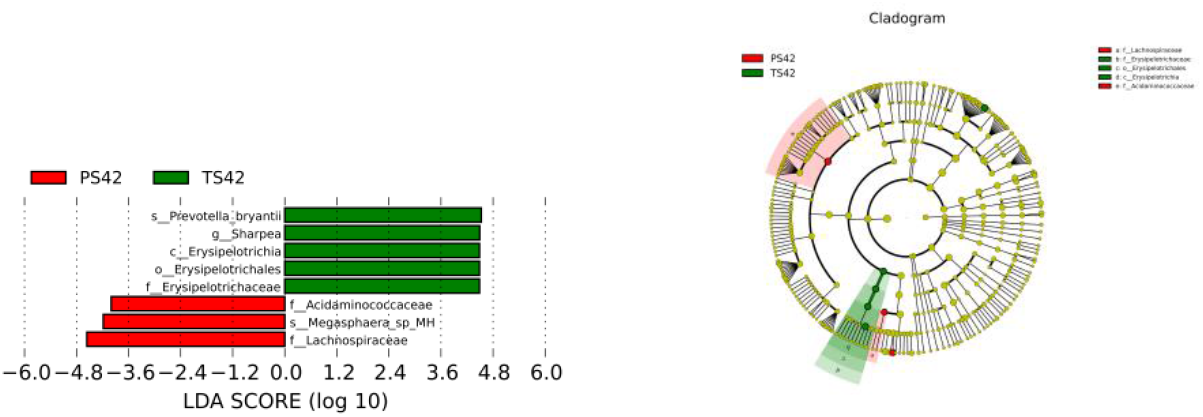
Value of linear discriminant analysis (LDA) effect size (LEf Se) on rumen microflora between two starter feeds for post-weaned lambs

### Functional prediction of rumen microbe

Functional profiles (KEGG level 2 pathways) of the two groups for pre-weaning and post-weaning lambs were all found to be similar in comparison (Fig 8). And the main functions of rumen microbe (the top seven) for lambs were replication and repair (pre-weaning: PS 10.56%, TS 11.03%; post-weaning: PS 11.44%, TS 11.39%), carbohydrate metabolism (pre-weaning: PS 11.04%, TS 10.95%; post-weaning: PS 10.73%, TS 10.72%), translation (pre-weaning: PS 9.86%, TS 10.33%; post-weaning: PS 10.65%, TS 10.67%), membrane transport (pre-weaning: PS 10.15%, TS 8.57%; post-weaning: PS 8.36%, TS 8.29%), amino acid metabolism (pre-weaning: PS 8.35%, TS 8.31%; post-weaning: PS 8.16%, TS 8.22%), nucleotide metabolism (pre-weaning: PS 4.66%, TS 4.84%; post-weaning: PS 5.00%, TS 4.99%), energy metabolism (pre-weaning: PS 4.26%, TS 4.55%; post-weaning: PS 4.60%, TS 4.65%). Only the “Transporters” and “Fatty acid degradation” predicted function of KEGG level 3 pathways of PS were increased significantly and “Amino acid related enzymes” was decreased significantly for pre-weaning lambs compared to those of TS (*P* < 0.05, Fig 9). And the predicted function of KEGG level 3 pathways between PS and TS for post-weaning lambs still had no significant differences (*P*>0.05).

**Fig 8.**
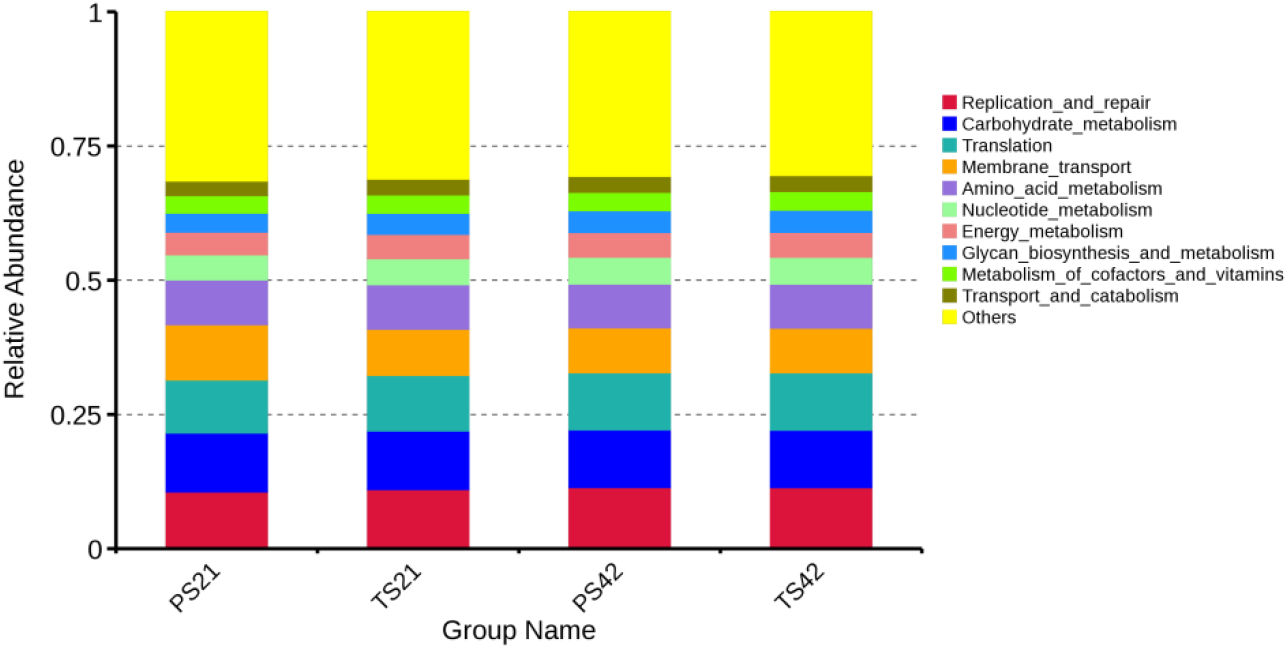
Relative abundances of top 10 function of rumen microflora between two starter feeds for pre-weaning and post-weaned lambs (KEGG level 2 pathways)

**Fig 9.**
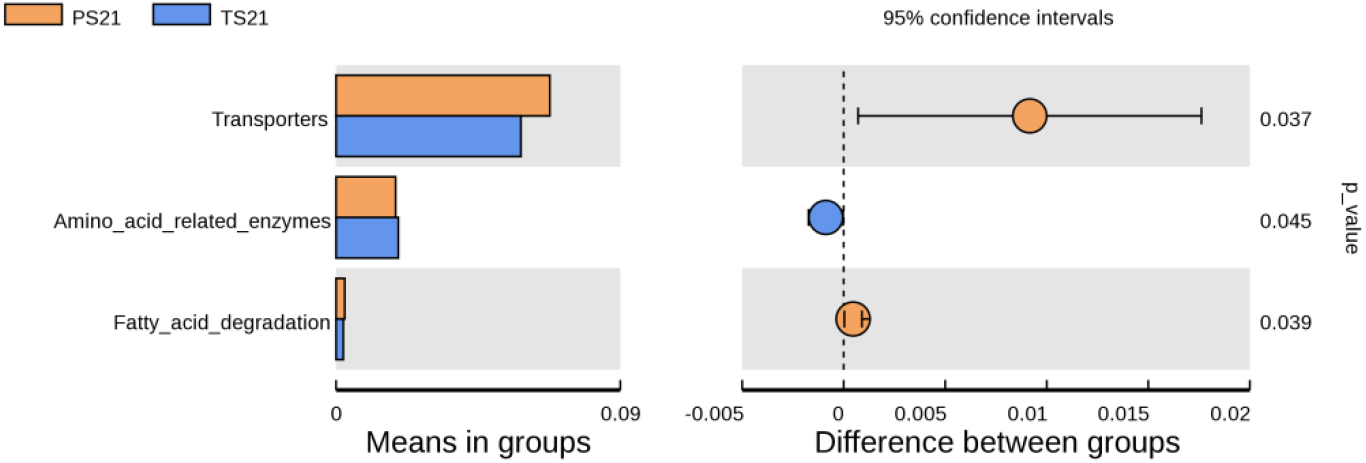
Analysis of different metabolism pathways between two starter feeds for pre-weaning lambs (KEGG level 3 pathways)

## DISCUSSION

Rumen pH was the most intuitive indicators reflected the fermentative condition of rumen, which could integrated reflect the rumen microbe, production, absorption and neutralization of organic acid. Furthermore, the acidity of rumen played a dominant role to maintain ruminal environment. Many factors affected the rumen pH, such as diet structure, secretion volume of saliva, speed of intake, VFA in rumen and the rates of production, absorption and excretion of organic acids. But the fundamental reasons of fluctuation of rumen pH were the diet structures and nutrition levels. Murphy and Kennelly indicated that rumen pH changed regularly from 5.0 to 7.5, which resulted from diet nature and time after intake (15). When intake fermentable carbohydrate, a lot of VFA could be produced that led to decrease of pH in rumen. In experiment, rumen pH changed from 5.14 to 5.96, which were in normal scope. However, rumen pH of TS was lower than them of PS respectively for pre-weaned and post-weaned lambs. The possible reasons were TS contained steam-flaked corn, which included a lot of fermentable carbohydrate and reduced rumen pH. Nejad et al. showed steam-flaked corn increased the gelatinization of corn starch so as to change the degradation form of corn in rumen microbe, increase the surface area of corn and enhance hydrolytic ability of rumen microbe and enzyme to starch granule (16). Hence, TS increased fermentable carbohydrate utilized by rumen microbe, which produced a lot of VFA and reduced the rumen pH.

VFA were main products fermented by carbohydrates in rumen, included the acetate, propionate and butyrate, which were important energy sources of ruminant. Research found VFA were important promoting factors to growth and development of rumen epithelium. Furthermore, among VFA, stimulant action of butyrate was the most effective, following as propionate and acetate (17). In experiment, concentration of total VFA, propionate and butyrate of TS were all higher significantly than them of PS for pre-weaned and post-weaned lambs, which showed TS were better benefits to fermentation and development of rumen. These were consistent with the results of research about calves. Lesmeister et al. reported concentrations of VFA and propionate of TS contained steam flaking corn were higher than them of starter feed contained the whole corn, dry-rolled corn and roasted-rolled corn in serum of calve (18). Pavlata et al. also found concentrations of VFA, acetate and propionate of TS with chopped straw were higher significantly than them of PS (19). These results indicated that texture starter feed contained steam flaking corn could provide more chemical stimulations to development of rumen, which were better benefits to fermentation and development of rumen for lambs. These were consistent with results of rumen development in experiment.

NH_3_-N was degradation products of protein nitrogen, non-protein nitrogen of diet and endogenous nitrogen in rumen, which were essential precursors to synthetize bacterial protein of rumen microbe. Maintaining reasonable NH_3_-N concentration was an important condition to growth and reproduction of rumen microbe. NH_3_-N concentration not only could reflect the speed of production and utilization, which degraded from nitrogenous substances by rumen microbe, but also could reflect the balances between degradation and synthesis of protein under specific diets to the extent. Many factors could affect the NH_3_-N concentration in rumen, such as protein quality of diets, emptying rate of chyme and absorption of rumen wall (20). It was reported that 5 mg/100ml NH_3_-N concentration was the lowest concentration which maintained growth and protein synthesis of rumen microbe. When NH_3_-N concentration was lower than 5 mg/100ml, the growth of rumen microbe would be suppressed. Moreover, the optimum NH_3_-N concentration which maintained growth of rumen microbe was 6.3-27.5mg/100m1 (21). In current experiment, the starter feed of two different physical forms did not affect the NH_3_-N concentration of rumen for pre-weaning and post-weaning lambs, but the NH_3_-N concentration of rumen were under the optimum scopes maintained growth of rumen microbe. Beharka et al. and Pazoki et al. found that the NH_3_-N concentration in rumen liquid of calves had no significant differences between pelleted and textured starter feed (22, 23). Additionally, Qi and Ga et al. also found that the corn processed by pelletizing, baking and steaming did not affect significantly the NH_3_-N concentration in rumen liquid of sheep (24, 25).

OTU could reflect the clustering quantities, and alpha diversities were used to evaluate the abundances and diversities of rumen microbe. Among alpha diversities, chao1 and Ace reflected the abundances of rumen microbe, whose higher chao1 and Ace indicated abundances of flora were greater; shannon and simpson reflected the diversities of rumen microbe, whose greater shannon and lower simpson showed diversities of flora were more (26). In experiment, OTU, chao1, ACE, shannon and simpson of rumen microbe all had no significant differences between two starter feed of different physical forms for pre-weaned and post-weaned lambs, only OTU of rumen microbe of TS had trends to lower than that of PS for pre-weaned lambs. These showed physical forms of starter did not affect the species, abundances and diversities of rumen microbe for pre-weaning and post-weaning lambs, only the species of rumen microbe of TS had trends to lower than that of PS for pre-weaning lambs. The results showed the lambs were easier to adopt the TS during courses of transition from liquid to solid starter feed, and could urge rumen of lambs to establish dominant flora and disappear instantaneous flora.

Intake of starter feed early could change rumen flora for pre-weaning lambs (27). With increasing of ages and intake of starter feed for lambs, *Bacteroidetes*, *Firmicutes* and *Proteobacteria* could become the main dominant flora of higher relative abundances, and the relative abundances of *Bacteroidetes* were increased, and the relative abundances of *Firmicutes* and *Proteobacteria* were reduced (2). When calves intake MR and starter feed, the *Proteobacteria* of rumen could be replaced by *Bacteroidetes* at ages of 42d, and the relative abundances of *Bacteroidetes* become the highest, which were possible to be related to chemical composition of diets (12). In experiment, the *Firmicutes* and *Proteobacteria* of TS had been replaced by *Bacteroidetes* for pre-weaning lambs, and *Bacteroidetes* had become the first predominant Phylum, and achieve similar relative abundances to them of TS for post-weaning lambs (pre-weaning, 61.96%; post-weaning, 65.36%). However, the *Firmicutes* and *Proteobacteria* of PS were replaced by *Bacteroidetes* for post-weaning lambs, and *Bacteroidetes* become the first predominant Phylum. Furthermore, relative abundances of *Bacteroidetes* of PS (57.28%) for post-weaning lambs were still lower than them of TS for pre-weaning (61.96%) and post-weaning (65.36%) lambs. The results showed the lambs were easier to adopt the TS during courses of transition from liquid to solid starter feed, and could urge rumen of lambs to establish dominant flora.

Among the rumen microflora of ruminants, *Bacteroidetes* and *Firmicutes* were two main dominant floras (28). It was well known that rumen microbe, especially *Bacteroidetes*, played an important role in degradation of starch and protein of diets, synthesizes of protein of rumen microbe, absorption of peptides and amino acids (29). And *Firmicutes* contained many bacterial degraded fibres, such as *Ruminococciis*, *Eubacterium*, *Pseudobutyrivibrio*, *Butyvibro* and *Oscillibacter*, whose main function were to degrad the cellulose (30). These showed *Bacteroidetes* was better benefits to degrade the concentrated diets, and *Firmicutes* was better benefits to degrade the roughage diets. Jiang et al. found the relative abundances of *Bacteroidetes* of fermented corn gluten meal were significant higher than them of corn gluten meal, and relative abundances of *Firmicutes* of fermented corn gluten meal were significant lower than them of corn gluten meal (31). In experiment, compared to PS, TS increased the relative abundances of *Bacteroidetes* for pre-weaning (PS, 35.74%; TS, 61.96%) and post-weaning (PS, 57.28%; TS, 65.36%) lambs, decreased the relative abundances of *Firmicutes* for pre-weaning (PS, 37.88%; TS, 32.08%) and post-weaning (PS, 31.21%; TS, 24.03%) lambs. These results were consistent with previous research. These results indicated physical forms of starter feeds affected the structures of rumen microbe at levels of phylum for pre-weaning and post-weaning lamb, and TS was better benefits to promote the fermentation of rumen and absorption of nutrients for lambs. The possible reasons were corns of TS processed by steam flaking, and improved the gelatinization of starch, contained more fermentable carbohydrates, which were better benefits to fermentation of *Bacteroidetes*. These were also consistent with results of fermentative parameters of rumen.

However, Kim et al. found the dominant flora in rumen was *Firmicutes* and *Bacteroidetes* in turn (32, 33). The reasons of differences might be related to composition of diets. Their diets were type of roughage, and the diets were type of concentration in current experiment. Additionally, the research also found when fed diets of higher proportional concentration, the dominant flora in rumen were *Bacteroidetes*; and when fed diets of higher proportional roughage, the dominant flora in rumen were *Firmicutes* (34). And *Bacteroidetes* were correlated negatively to *Firmicutes* (35). These proved further TS were better effective to fermentation of rumen for pre-weaning and post-weaning lambs.

The main fermented products of *sharpea* in rumen of sheep were lactates, and formation of lactates promoted further fermentation of *sharpea*, urged lactates to change into butyrate, which led to produce lower H_2_ compared to traditional fermentation directly from carbohydrate to butyrate and reduced production of CH_4_ in rumen (36, 37). In current experiment, compared to PS, TS increased significantly the relative abundances of *sharpea* of rumen for pre-weaning and post-weaning lambs, which were consistent with significant higher concentration of butyrate of TS in rumen liquid. These results showed TS contained steam flaking corn were benefits to improve rumen microflora, urge rumen fermentation for lambs. At same time, Xue et al. also found concentrations of butyrate in rumen liquid were correlated positively to the relative abundances of *sharpea* when they compared effects of higher yield, higher concentrations of milk protein and lower yield, lower concentrations of milk protein on rumen microflora of calves (38). Lin et al. also proved when fed starter feed, *Sharpea* produced lactates was main enriched in rumen of lambs (39).

Additionally, we found PS also increased significantly the relative abundances of *unidentified_Clostridiales*, *Lactpcoccus*, *Lactobacillus*, *Sarcina* and *unidentified_Cyanobacteria* for pre-weaning lambs compared TS in the experiment. *Clostridiales* and *Lactpcoccus* were all the main representative genus of *Firmicutes*, which could degrade cellulose in rumen. Hence, the higher relative abundances of *Clostridiales* and *Lactpcoccus* of PS were consistent with higher relative abundances of *Firmicutes* of PS (PS, 37.88%; TS, 32.08%) at levels of phylum for pre-weaning lambs. *Lactobacillus* could ferment carbohydrates to produce lactates, which were benefits to health of animals. Hence, *Clostridiales*, *Lactpcoccus* and *Lactobacillus* played decisive roles in digestion and absorption of nutrients in gastrointestinal tract and immunities of animals. *Cyanobacteria* were one of microalgaes, which could improve the performances of animals and qualities of meat as feed additives (40). *Sarcina* was related to rumen bloating of lambs (41) and calves (42), which should not exist in digestive tracts of animals.

However, predicted functions of rumen were found to be similar in two groups for pre-weaning and post-weaning lambs in current experiment. Only the “Transporters” and “Fatty acid degradation” of PS were increased significantly and “Amino acid related enzymes” was decreased significantly for pre-weaning lambs compared to those of TS. It was confirmed previously that significant changes of microbial composition might not lead to a shift of function because many microbes shared the same metabolic pathways. Li et al. found that all of the functional classes between two age groups (d14 and d42 of calves) were similar, suggesting that although their phylogenetic composition greatly fluctuated, the rumen microbial communities of pre-ruminant calves maintained a stable function and metabolic potentials (12). These might be the reasons that two group lambs had similar functions in this experiment. Maybe it was necessary to analyse the rumen microbiome functions using metagenomic and/or metabolomics technologies for completed and integrated understand the impact of rumen function further.

In a word, physical forms of starter feeds affected the fermentation and microbial composition of rumen for pre-weaning and post-weaning lambs. TS were better benefits to improve fermentation environment and establish dominant flora of rumen early, which were in favour of growth and development of rumen for pre-weaned and post-weaning lambs.

## MATERIALS AND METHODS

### Animals, feeds and experimental design

This experiment was carried out on a local sheep farm (Baiyin Kangrui breeding sheep co., Baiyin, Gansu province). All the experimental protocols performed in this study were approved by the Animal Care Committee of Gansu Agricultural University and the experimental procedures used in this study were in accordance with the recommendations of the University’s guidelines for animal research. In this study, twenty-four healthy male Hu lambs, whose average body weight were 5.04±0.75 kg, were separated from their dams at day 8 and moved into a naturally ventilated barn with individual cages (0.8 ×1.3 m). And the trial lambs were fed either milk replacer (MR) and pelleted starter feed (PS, with a mean particle size of six mm diameter), or MR and textured starter feed (TS, which included coarse mashed steam-flaked corn, also with a mean particle size of six mm diameter) in pre-weaning (day 8 to 35) and post-weaning (day 36 to 42) lambs. And the MR (23% CP and 12% fat, DM basis) was fed by single bottles to lambs at 2% of body weight at day 8 divided as three equal amounts at 08:00, 14:00 and 20:00 in pre-weaning. After weaning, all lambs continued to be fed starter according to their trial group. And all lambs had free access to readily avail clean fresh water and their respective ad lib starter feed in the whole experiment. The diets (Table 3) were prepared by Gansu Aonong Feed Co., Ltd according to National Research Council recommendations (NRC, 2007). The MR was bought from Beijing Accurate Animal Nutrition Research Center.

**Table 3.** Ingredients and chemical composition of the experimental diets.

| Item | Pelleted | Textured |
| --- | --- | --- |
| Ingredients (g/kg) |  |  |
| Alfalfa hay | 50.00 | 50.00 |
| Corn grain, ground | 650.00 | - |
| Corn grain, steam flaked | - | 650.00 |
| Wheat bran | 50.00 | 50.00 |
| Expanded soybean | 60.00 | 60.00 |
| Soybean meal | 165.00 | 165.00 |
| Salt | 3.00 | 3.00 |
| Calcium carbonate | 11.80 | 11.80 |
| Vitamin and mineral mix <sup>a</sup> | 10.00 | 10.00 |
| Sweetening agent | 0.20 | 0.20 |
| Total | 1000.00 | 1000.00 |
| Chemical composition (g/kg, DM) |  |  |
| Apparent digestible energy b(MJ/kg) | 13.75 | 13.82 |
| Dry Matter <sup>b</sup> | 892.40 | 885.00 |
| Crude Protein <sup>b</sup> | 214.60 | 216.40 |
| Ether extract <sup>b</sup> | 27.20 | 27.90 |
| Neutral detergent fiber <sup>b</sup> | 128.00 | 128.80 |
| Non-fiber carbohydrates <sup>c</sup> | 625.10 | 622.20 |
| Starch <sup>b</sup> | 115.10 | 117.10 |
| Calcium <sup>b</sup> | 7.50 | 7.10 |
| Total phosphorus <sup>b</sup> | 4.48 | 4.52 |

<sup>a</sup> Provided per kilogram of premix: Fe 75000 mg, Zn 15000 mg, Cu 3500 mg, Mn 15000 mg, I
500000 mg, Se 50 mg, Co 200 mg, VA 2500000 IU, VD 1000000 IU, VE 1900 IU.
<sup>b</sup> The actual values for measurement.638 <sup>c</sup> Non-fiber carbohydrates=1000 – (crude protein + ether extract + aNDF + ash) (NRC, 2001).

### Sample collection

Lambs for each treatment were euthanized by captive bolt stunning and exsanguinated in a specialized room of the experimental farm without any transportation at the age of 21 or 42 days. After slaughter, a part of rumen content was used to collect rumen liquid by immediately filtering through four layers of swab and transferring into 10 cm centrifuge tubes, and then stored at −20°C to analyse the total volatile fatty acids (VFAs) and ammonia nitrogen (NH_3_-N); the other part of rumen content was collected for storage at − 80 °C for rumen bacteria analysis.

### Determination of rumen fermentative parameters

After slaughter, the rumen content was mixed and determined pH immediately by PB-10 acidity meter (Zedorius, Kogentin, Germany). Rumen liquid were thawed and analyzed for individual and total VFA concentrations by gas chromatography (AI 3000, Thermo, Germany) (43) and NH_3_-N by colorimetric method (44) using visible spectrophotometer (Shanghai Jinghua Technology Co. Ltd). Details of VFA determination were as follows:

0.6μL rumen liquid samples were injected by an auto sampler into an AE-FFAP (30m× 0.25mm× 0.33 μm; Zhongke Antai, Lanzhou, China). Chromatographic conditions: temperature of injection entrance 200°C; N_2_ flow 2.0 mL•min^−1^; split ratio 40:1; procedure heating mode (120°C 3 min, 10°C•min^−1^ to 180°C, kept 1 min); detector temperature 250°C; FID air, H_2_ and N_2_ flow were 450 mL•min^−1^, 40 mL•min^−1^ and 45 mL•min^−1^ respectively; cylinder heating procedure: from 45°C to 150°C as speed 20°C• min^−1^, and kept 5min. Finally, the peak integration was performed using Chromeleon^®^ Software.

### Total DNA extraction of rumen microbe, illumina sequencing

Rumen content was sent to Novogene Bioinformatics Technology Co., Ltd. (Beijing, China) to extract DNA and sequence of rumen microbe. Details were following:

Total DNA of rumen microbe in rumen content was extracted by thecetyltrimethylammonium bromide method (45) with a bead-beater (Biospec Products; Bartlesville, OK, United States) as described by Gagen et al. (46). The amplification of V_3_-V_4_ hypervariable region of the 16S rRNA gene was carried out with formwork of each of the DNA samples using the primer set 515F/806R and Phusion^®^ High-Fidelity PCR Master Mix (New England Biolabs, Ipswich, MA, United States) as described by Caporaso et al. (47). When each forward and reverse primer had a 6-bp error-correcting barcode at the 5’ terminus, it was seen as unique to each DNA sample. The sequencing for all samples was on an Illumina HiSeq platform by Novogene Bioinformatics Technology Co., Ltd. (Beijing, China) to generate 2 × 250 bp paired end reads.

After the paired-end reads were cut off barcode and primer, they were joined together and formed single sequences using FLASH based on overlapping regions (48). Sequences with a quality score of <20 and a length of >300 bp or <200 bp were filtered and discarded using Quantitative Insight into Microbial Ecology (QIIME, 49). At same time, the possible chimeric sequences were also identified and removed from the sequences using the usearch61 algorithm in USEARCH 6.1 (50). Operational taxonomic units (OTUs) were clustered as 97% similarity, and chosen the representative sequence according to the algorithmic principle and annotation analysed in SILVA SSU rRNA datebase (51) used Uparse (Uparse v7.0.1001, 52) by Mothur method (53).

### Microbial functional prediction

Microbial function predicted byTax4Fun based on 16S Silva database (54). Details were as following:

All the genes 16S rRNA sequences of prokaryote in Kyoto Encyclopedia Genes and Genomes (KEGG) database were extracted, then compared in SILVA SSU Ref NR database (BLAST bitscore >1500)by BLASTN method and established correlation matrix. Finally, functional information of all the genes 16S rRNA sequences of prokaryote annotated in KEGG database were compared with functional information in SILVA database by UProC and PAUDA so as to achieve the goals of microbial functional prediction.

### Statistical analyses

Data for rumen fermentative and metabolic parameters, alpha diversity indices (number of OTU, ACE, Chao1, Shannon and Simpson index) of rumen microbe were pooled at each time point for the six slaughtered lambs in each group. Data were analysed as independent sample t-tests (SPSS 20.0, Inc., Chicago, IL, USA). Significance was designated as *P*<0.05 with a trend being between *P*≥0.05 and *P*<0.10. Beta diversity of rumen microbe was analysed by T-test and wilcox-test. Analysis of similarity between groups was calculated by Bray-Curtis. Principal co-ordinates analysis (PCoA) of rumen bacterial OTUs calculated the distance by Bray-Curtis firstly, then drawn the fig by R software (v3.3.0). Difference between groups was analysed by LEfSe (LDA Effect Size), and LDA>4 was different marking of statistics and biology (55).

## ACKNOWLEDGMENTS

This work was supported by National Natural Science Foundation of China (31860654). The authors express their kind appreciation to Dr. Xiaojuan Wang and Ligang Yuan from the Gansu Agricultural University for their assistance throughout the experiments.

